# Using the structure of genome data in the design of deep neural networks for predicting amyotrophic lateral sclerosis from genotype

**DOI:** 10.1101/533679

**Authors:** Bojian Yin, Marleen Balvert, Rick A. A. van der Spek, Bas E. Dutilh, Sander Bohté, Jan Veldink, Alexander Schönhuth

## Abstract

Amyotrophic lateral sclerosis (ALS) is a neurodegenerative disease caused by aberrations in the genome. While several disease-causing variants have been identified, a major part of heritability remains unexplained. ALS is believed to have a complex genetic basis where nonadditive combinations of variants constitute disease, which cannot be picked up using the linear models employed in classical genotype-phenotype association studies. Deep learning on the other hand is highly promising for identifying such complex relations. We therefore developed a deep-learning based approach for the classification of ALS patients versus healthy individuals from the Dutch cohort of the ProjectMinE dataset. Based on recent insight that regulatory regions on the genome play a major role in ALS, we employ a two-step approach: first promoter regions that are likely associated to ALS are identified, and second individuals are classified based on their genotype in the selected genomic regions. Both steps employ a deep convolutional neural network. The network architecture accounts for the structure of genome data by applying convolution only to parts of the data where this makes sense from a genomics perspective.

Our approach identifies potential ALS-associated genetic variants, and generally outperforms other classification methods. Test results support the hypothesis that ALS is caused by non-additive combinations of variants. Our method can be applied to large-scale whole genome data. We consider this a first step towards genotype-phenotype association with deep learning that is tailored to genomics and can deal with genome-sized data.

## 1 Introduction

Amyotrophic lateral sclerosis (ALS) is a neurodegenerative disease affecting the upper and lower motor neurons, resulting in a progressive loss of muscle strength leading to paralysis and eventually death (Goldstein and Abrahams, 2013; Phukan *et al.*, 2007). For many patients ALS is likely caused by genetic aberrations. While a handful of major genetic risk factors have been identified, no more than 15% of the heritability has been explained so far (Van Rheenen *et al.*, 2016). This is because the genetic architecture of ALS has been found to be rather involved: ALS seems to be evoked through not necessarily additive combinations of genetic aberrations that individually only have a small effect and can thus not be detected using the currently available genotype-phenotype association approaches (Van Rheenen *et al.*, 2016).

Motivated by these findings, the application of prediction and/or association schemes that can capture non-additive effects is very promising. More than that, the evaluation of more complex schemes might even be an urgent necessity if one aims at further progress in predicting ALS, associate it with genetic causes, and, eventually, also treat it successfully.

In the last ten years, the identification of genotype-disease relations has been considerably enhanced by the use of large-scale genome data. Project MinE is an international initiative to collect genome data of tens of thousands of ALS patients and healthy control individuals. Many individuals have been sequenced at considerable depth of genome coverage (Project MinE ALS Sequencing Consortium and others, 2018). The corresponding wealth of data is still awaiting its full exploration. Clearly, it carries the potential for pointing out ALS risk fators, guiding further research and drug development.

Genome-wide association studies (GWAS) are the current state-of-the-art in analyzing genotype-phenotype data. Statistical tests are used to determine the level of association between a single genetic variant and phenotype, and are therefore suitable for uncovering genotype-phenotype associations that involve single variants or variants interacting with others in additive schemes. GWAS have successfully identified disease-associated variants over a wide range of disorders (Visscher *et al.*, 2017) including ALS (van Es *et al.*, 2009; Nicolas *et al.*, 2018). However, the approach has been found to be unable to find the non-additive combinations that are associated with phenotypes (Wray *et al.*, 2013), which limits its power as genetic variants often constitute phenotype in non-additive combinations. This could for example be caused by epistasis, where the effect of one variant on phenotype is dependent on the presence or absence of others (Frankel and Schork, 1996; Moore, 2003). As above-mentioned, the genetics underlying ALS have been found to be more involved and are therefore unlikely to be fully unravelled using basic association schemes (Van Rheenen *et al.*, 2016). The application of novel data analysis approaches that account for complex interactions between genotype input variables and ALS are thus very promising.

Thanks to advances in the recent past, deep neural networks (DNNs) have turned into powerful classifiers in several application areas including bioinformatics (Angermueller *et al.*, 2016). They have been proven to map arbitrarily complex relationships between multiple input features (in our case genetic variants) and output labels (here for example binary-valued labels ‘ALS’ or ‘no ALS’). In addition, DNNs have been pointed out to be particularly big data compatible (Schmidhuber, 2015). That is, they can handle a considerably larger number of input variables than most other machine learning methods, a prerequisite for the analysis of genome data. DNNs therefore hold the clear promise to successfully map complex genotype-phenotype associations.

DNNs cannot, however, be applied off-the-shelf when mapping genetic variants to disease (ALS) status; several hurdles need to be overcome. The first is the size of genome data: the sheer number of input variables (genetic variants, which amount to usually millions) exceeds the number that these models can deal with easily (a few hundred of thousands). Second, while DNNs can achieve great classification accuracy, interpretability is insufficient: it is difficult to determine why a DNN classified a sample as a case or a control. This is a major drawback for genotype-phenotype association studies, as the main goal is to identify (combinations of) variants that associate with disease rather than obtaining a high classification accuracy. Third, DNNs have delivered their most striking successes when applied in image classification tasks. High classification accuracies were obtained with networks of great depth, employing the hierarchical nature of these images (pixels together form lines, which together form basic shapes, etcetera). Thereby, the employment of convolutional filters/layers have been crucial in delivering the breakthroughs. Such filters make use of the position invariance of local structures in images, a property that does not hold for genome data.

A few studies have considered using deep learning for genotype-phenotype association studies. Most approaches first reduced the number of variants included in the model either by selecting variants that were known to be associated with disease (Uppu and Krishna, 2017; Hess *et al.*, 2017), or by preselecting those variants that showed a sufficiently strong correlation with phenotype in a regular GWAS (Montañez *et al.*, 2018b; Bellot *et al.*, 2018). Two studies combine the latter strategy with the use of autoencoders for further dimensionality reduction (Montañez *et al.*, 2018a; Fergus *et al.*, 2018). These approaches have the same drawback as a classical GWAS: already in the preselection step epistasis is overlooked, and variants that have a small effect on their own will not be included in further analysis. An alternative approach is proposed by Romero *et al.* (2016). The authors limit the computational burden by considering the transpose of the data matrix, which is similar to considering features as samples and vice versa, to learn the model parameters. As the number of genetic features is much larger than the number of samples in a genotype-phenotype association study, this leads to a major reduction in the number of trainable parameters and hence strongly reduces the time required for training. Tran and Blei (2017) define an implicit causal model that aims to identify relations between variants, and deals with the data dimensionality by updating the model one variant at the time. In summary, while a couple of earlier studies have used deep learning to predict phenotype from genotype, only two were able to deal with several hundreds of thousands of genetic variants. None have employed the structure inherent to genome data, and interpretation of the results has not been addressed in these studies.

This paper presents novel deep neural network architectures and a protocol by which to predict the occurrence of ALS from individual genotype data. In summary, we developed a deep learning-based method that (1) can handle genome-sized data by pre-selecting parts of the genome that are most relevant for classification, (2) provides insight in which genomic regions are relevant to classification, and (3) is capable of classifying ALS patients versus healthy control individuals from genome data. The design of our approach in general and our network architecture in particular is driven by the structure of genome data.

We demonstrate in our experiments that by means of our new architectures, we achieve more than 75% accuracy in predicting ALS from genotype data when considering chromosomes 7, 9, 17 and 22. Our results demonstrate that our ALS-Net clearly outperforms other machine learning tools and protocols we have been experimenting with, and drastically outperform state-of-the-art GWAS style prediction technology that is based on logistic regression. Our results therefore demonstrate that prior knowledge on the structure of genome data can aid in the design of a deep learning-based approach and the neural network architectures to yield improved accuracy rates in classifying genotypes with respect to occurrence of ALS. At the same time, we are aware that here we have only made the first steps towards routine application of deep neural networks in classifying genetically involved diseases from individual genomic profiles. We will point out where further improvements are conceivable along the way in the following, convinced that we are, at the very least, providing a very promising template for further explorations along this avenue of research.

## 2 Approach

We propose to make use of prior knowledge to tackle the dimensionality issue inherent to working with genome-sized data. The majority of the millions of variants in genome data are irrelevant, as these are not involved in disease. It has been found in general that most variants that relate to disease phenotypes reside in the DNAse hypersensitive sites (Maurano *et al.*, 2012), that is, in the majority of cases they occupy the promoter regions preceding genes, where transcription is initiated. We therefore focus on the promoter regions.

An interpretable model is able to indicate which genomic regions were relevant to classification. We therefore developed a two-step approach to employ neural network architectures for mapping associations between genotypes and the occurrence of ALS. The first step consists of individual classifiers for each promoter region, i.e., individuals are classified based on their genomic information from a single promoter region only. The classification accuracy obtained with an individual promoter region is an indication for the region’s predictive power, and only the eight best performing promoter regions are considered for further analysis. In the second step the genome information of the selected promoter regions is combined and an overall classifier is trained for final classification. This is illustrated in Figure 1, where we denote the promoter region-specific neural network by Promoter-CNN (CNN for convolution neural net) and the network that classifies samples based on a combination of promoter regions by ALS-Net. We develop and validate our approach using GWAS data from the Dutch cohort of ProjectMinE, which contains 4,511 cases and 7,397 controls.

**Figure 1:**
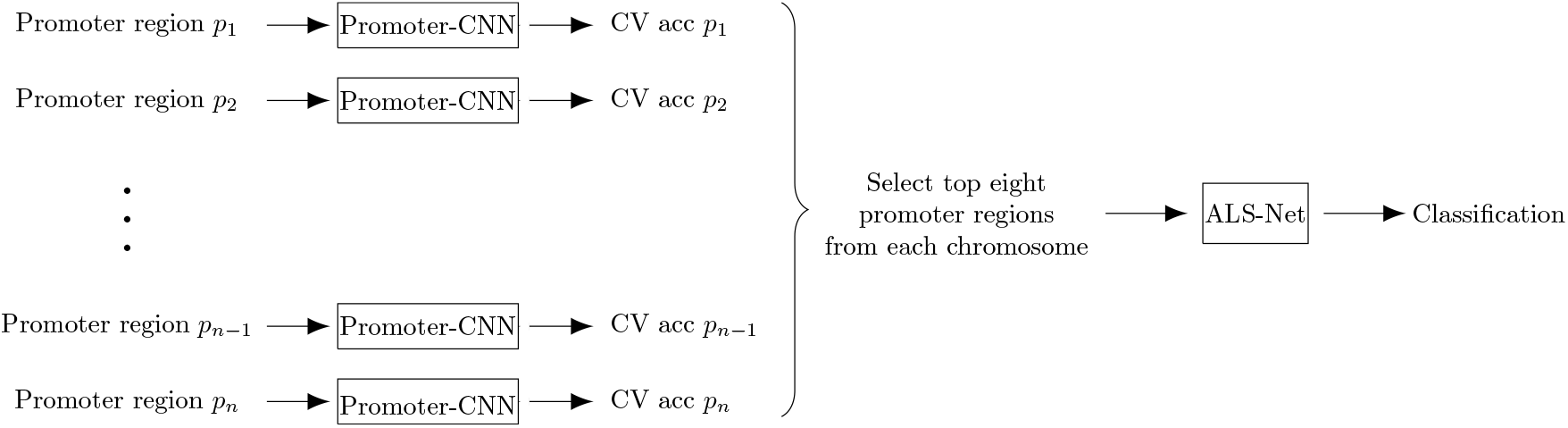
An overview of the workflow. CV = cross validation, acc = accuracy.

As noted before, the success of DNNs for image classification heavily relies on the local structures that are present in images. Genotype data do not convey neighborhood structures that are as easy to grasp as in images and applying convolution is less straightforward. Still, genome data does have a neighborhood structure which is due to two aspects. First, the genome consists of blocks that together form functional units, such as genes and promoter regions. Second, genetic variants are passed on from ancestor to offspring in terms of blocks rather than in isolation. Although in many cases details have not been fully understood, usually combinations of neighboring variants (haplotype blocks) are responsible for the establishment of phenotypes, rather than variants in isolation. This justifies the application of DNNs that take neighborhood structures into account (Bellot *et al.*, 2018). Note that the above does not contradict that isolated variants can be indicative of phenotypes: single variants usually are in linkage disequilibrium with other variants in their block, which establishes that basic GWAS can nevertheless be successful.

## 3 Methods

### 3.1 ProjectMinE data

We use data collected by ProjectMinE, a worldwide effort to collect whole-genome data from both ALS patients and unaffected individuals for the identification of ALS-causing variants (Project MinE ALS Sequencing Consortium and others, 2018). The dataset we used contains solely the Dutch cohort, consisting of 4,511 ALS patients and 7,397 healthy individuals. including 6,127 males and 5,781 females.

First SNPs were annotated according to dbSNP137 and mapped to the hg19 reference genome. Quality control (QC) was first performed per cohort to remove low quality SNPs and individuals using PLINK 1.9 (Purcell and Chang, 2015; Chang *et al.*, 2015) (--geno 0.1 and --mind 0.1). HapMap3 (Consortium *et al.*, 2010) projected PCs were calculated and extreme CEU population outliers were removed (25 SD). Cohorts were merged into strata based on genotyping platform. Subsequently, more stringent SNP QC was performed (--maf 0.01, --mind 0.02, --hwe 1e^−5^ midp include-nonctrl, --test-mishap excluded *p*¡1e-8) followed by more stringent individual QC (--geno 0.02, --het excluded ¿0.2, and removed sexcheck failures and missing phenotypes). We then only kept the autosomal regions. We filtered SNPs based on differential missingness (--test-missing midp) and excluded those with a *p*-value below 1e–4. Finally, we removed duplicated individuals (PI_HAT¿0.8) and filtered more stringently on population outliers (hm3: 10SD, 1KG: 4SD, Stratum itself: 4SD). Strata were then imputed using the HRC reference panel (Das *et al.*, 2016).

Motivated by the fact that chromosomes 7, 9 and 17 all have been found to carry elevated amounts of missing heritability (Van Rheenen *et al.*, 2016), we focus on those chromosomes. Additionally, we included chromosome 22 that was reported to have a low level of heritability. The genome data of the four chromosomes contains 823,504 positions of variation.

Note that all chromosomes occur in pairs: one maternal and one paternal copy. We convert the data, which is in VCF format, to minor allele frequency data. Hence the data of each individual is a list of values in {0, 1, 2}, indicating the number of occurrences of the minor allele at each position position on the genome. In some cases information for one of the chromosome copies is missing, and we assume this to be the frequent allele.

We focus on the promoter regions. As the position of a promoter region on the genome is generally not as well defined as the transcription start sites of a gene, and because a deep neural network requires the data representation for each promoter region to be of the same size, we used the following approach for determining the variants that are in the promoter regions. We used the transcription start sites as reported in the RefSeq database (O’Leary *et al.*, 2015). The 56 variant positions upstream and the 8 variant positions downstream of the transcription start site were then included in our representation of the promoter region. Hence, each promoter region is represented by a list of 64 values from the set {0, 1, 2}. Note that a gene can have multiple transcription start sites, and hence multiple promoter regions.

In summary, the input data to our model for one individual is a list of vectors in {0, 1, 2}^64^, where each vector contains the minor allele frequencies of a promoter region.

### 3.2 Neural network architectures

Promoter-CNN uses two convolution layers followed by two dense layers. As such (unlike ALS-Net in the following), Promoter-CNN is not deep, which is justified by the small input. Details of the architecture of Promoter-CNN are presented in Table 1. Batch normalization is applied after each layer, followed by the softplus activation function.

**Table 1:** Network architecture of the classifier with input data from a single promoter region. The output shape is given as (width, channels). BN = batch normalization, Act = softplus activation.

| Layer type | Description | Output shape |
| --- | --- | --- |
| Input |  | (64, 1) |
| Convolution, BN and Act | $1 \times 1$ filter,<br>4 output channels | (64, 4) |
| Convolution, BN and Act | $4 \times 4$ filter,<br>32 output channels | (61, 32) |
| Reshape | Flatten | (1952, 1) |
| Dense, BN and Act |  | (148, 1) |
| Dense, BN and Act |  | (16, 1) |
| Output | Softmax | (2, 1) |

ALS-Net is a more involved neural network, where the design of the architecture is based on intuition guided by the structure of genome data. This will be further explained below. The full network architecture is shown in Figure 3, with details of Blocks 1 up to 4 in Figures 4 up to 7.

The input is formed by concatenating the vectors with genome information from the individual selected promoter regions to obtain one vector of length 64 × 8 × *c* = 512*c*, where 64 is the number of variants in a promoter region, 8 is the number of selected promoter regions per chromosome and *c* is the number of chromosomes included in the analysis.

In the first block of layers each promoter region is considered separately, that is, the information from different promoter regions is not yet combined. This allows the model to focus on obtaining a good representation of the individual promoter regions before combining their information. The first layer of Block 1 is a convolution layer with stride 64, which ensures that information from separate promoter regions is not combined, and kernel size 64, which implies that the information from each promoter region is processed as a whole (so no convolution within the promoter region). The layer has 256 output channels, hence 256 functions of the input values of a single promoter are trained and the information from a promoter region is now represented by 256 values (see convolution step in Figure 2). The second layer is a convolution layer with kernel length 1, stride 1 and 256 output channels, as proposed by Howard *et al.* (2017). This layer takes a linear recombination of the information from each single promoter region. It does so 256 times with different weights and biases, once for each output channel. The first two convolution layers do not have an activation function. The block is concluded with a batch normalization layer and a rectified linear unit activation function.

**Figure 2:**
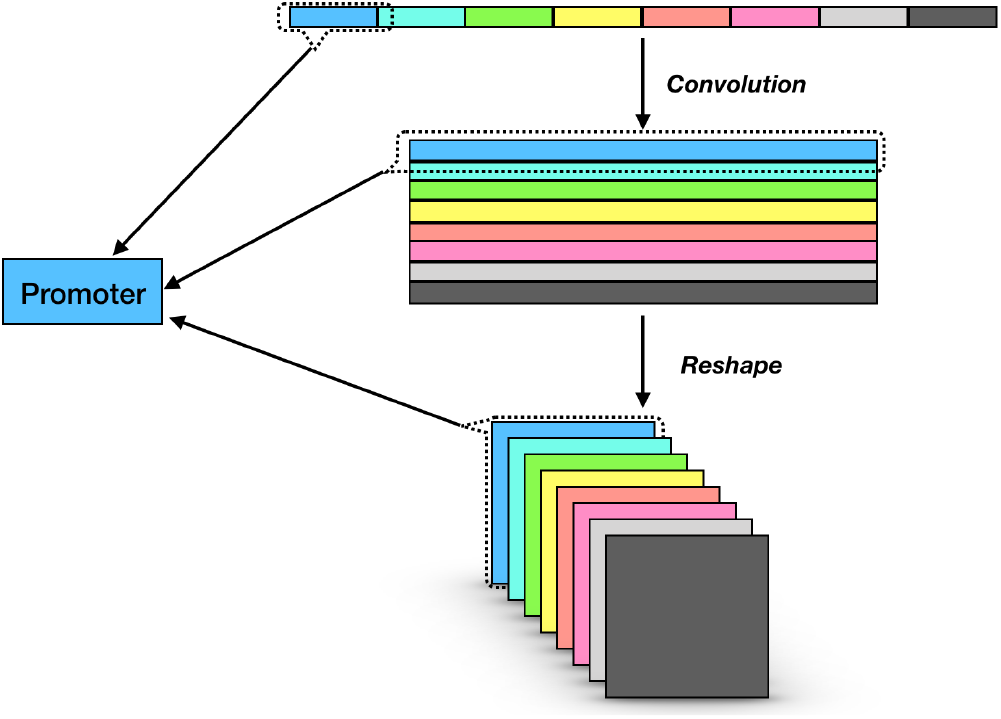
Change of tensor shape throughout Block 1 and Reshape.

Next, the tensor is reshaped into a 3 dimensional tensor, where the information of each promoter region is reshaped from a vector of length 256 to a 16 by 16 matrix (see reshape step in Figure 2). This three dimensional tensor can be viewed as an image of 16 by 16 pixels with 8*c* channels, where each channel corresponds to a promoter region.

In block 2 the promoter regions are combined. Since the underlying classification task requires the model to identify complex patterns we employ parallel computation blocks as well as residual connections to prevent the loss of information in future layers (Szegedy *et al.*, 2015; He *et al.*, 2016). Furthermore, several max pooling layers were included to prevent the model from overfitting. The model concludes with two dense layers. While dense layers are usually preceded by a flattening of the output of the previous layer, we make use of a global average pooling layer instead to allow for a strong dimensionality reduction.

### 3.3 Training and testing procedure

The dataset was split into a train-validate set (90% of samples) and a test set (10% of samples). The train-validate set was used for model development and selection of the promoter regions, the test dataset was used only for final testing. To test the model fairly, the ratio of cases and controls is 1:1 in the test dataset.

A nine-fold cross validation on the train-validate data was used to train Promoter-CNN. For each chromosome the eight promoter regions that achieved the best prediction accuracy averaged over the nine folds were selected for further analysis. The small network is trained using stochastic gradient descent on 50 epochs where the batch size is 64, and with a learning rate of 0.01.

The architecture and other hyperparameters for ALS-Net are optimized using a nine-fold crossvalidation of the train-validate data. The network architecture was optimized based on the learning curve and performance measures such as accuracy, precision and recall. Network parameters are optimized using the AdaGrad algorithm (Duchi *et al.*, 2011) with an initial learning rate of 0.02 and a decay of 2e^−4^. Optimization was performed over 300 epochs with a batch size of 32.

The model’s network architecture is optimized based on chromosome 7 only, and used for all four chromosomes individually as well as for the combination of the four chromosomes. Parameters are optimized separately for each chromosome as well as for the combined model.

The performance of our approach was tested by applying ALS-Net to the test data. Hence for these samples we only use the selected promoter regions.

### 3.4 Comparison with other machine learning approaches

The performance of ALS-Net is assessed using the test data, and is compared with the performance of logistic regression – this corresponds with the approach for calculating the polygenic risk score (PRS) (Dudbridge, 2013) –, support vector machine (SVM, Vapnik (1998); Joachims (1998)), random forest (Breiman, 2001) and AdaBoost (Friedman *et al.*, 2000; Freund *et al.*, 1999). For each of these we used the same promoter regions as for the large neural network. Hyperparameters were optimized using a cross validation approach, and performance on the test dataset is reported.

In logistic regression a linear function is used to estimate the disease risk score from genotype, followed by a classification where samples with a predicted risk score above a predetermined threshold are considered as positives (in our case ‘ALS’) and the others as negatives (in our case ‘no ALS’). While GWAS uses a single genetic variant as explanatory variable, we base our prediction of disease status on multiple variants, as is common for the calculation of the PRS. We apply logistic regression to the full set of promoter regions as well as to the variants that reside in the promoter regions selected by Promoter-CNN. Note that the choice of threshold determines the balance between precision and recall. In order to allow for comparison with the other methods, we chose the threshold such that accuracy on the training set is maximized.

SVM is a popular binary classification method designed to find a non-linear boundary (determined by the kernel function) to maximize the margin between two clusters. Here we used a radial basis function SVM with kernel coefficient 0.001.

Random forest is a widely used machine learning algorithm that creates multiple decision trees and combines their individual classifications to abstract a final classification. Using a large number of decision trees is required for higher accuracy, but also results in slow training. We implement a random forest consisting of 100 trees with maximum depth 5, and at most 100 features will be considered when looking for the best split.

The core idea of AdaBoost is to train several decision trees, assign weights to samples and classifiers to force the algorithm to focus on hard-to-classify samples, and combine the weighted classifications to form a stronger final classifier. While the model is powerful and yields explainable results, it is sensitive to outliers. We used AdaBoost with 1000 decision trees of depth 3.

**Figure 3:**
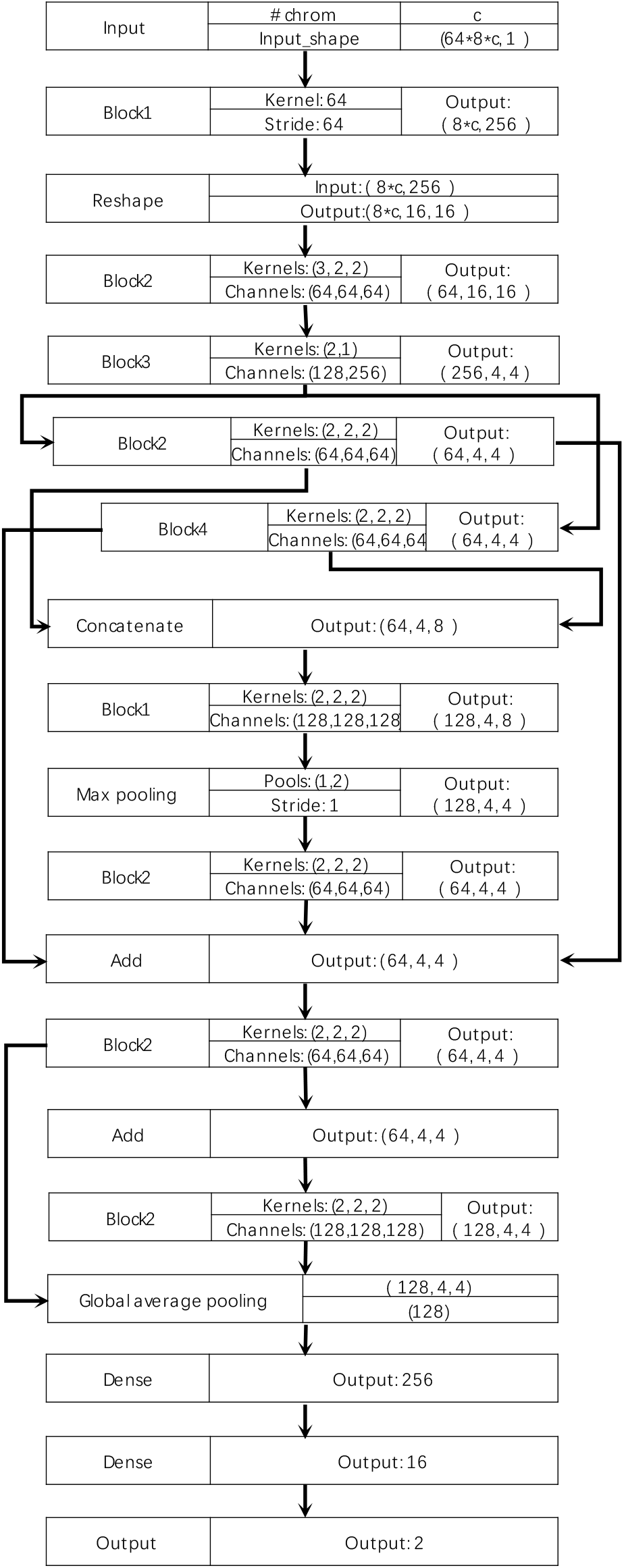
Architecture of ALS-Net. The details of Block 1, Block 2, Block 3 and Block 4 are provided in Figures 4, 5, 6 and 7, respectively.

**Figure 4:**
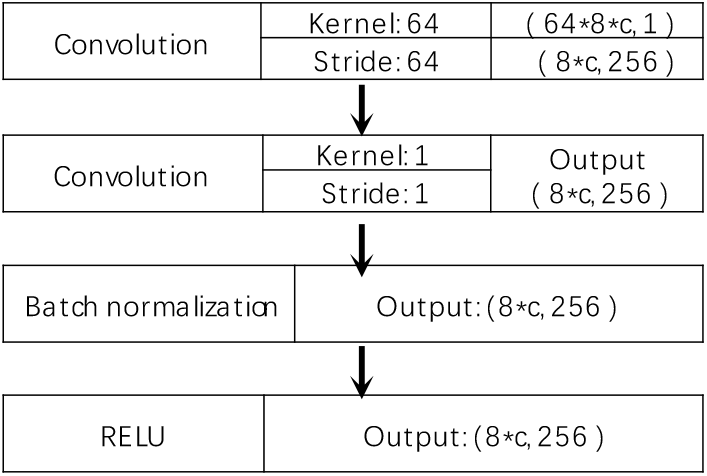
Network architecture of Block 1. *c* is the number of chromosomes included in the model.

**Figure 5:**
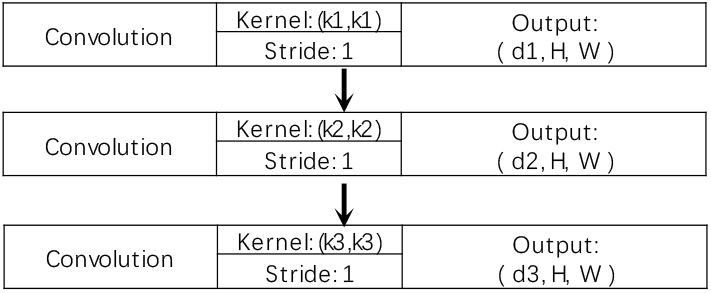
Network architecture of Block 2. Inputs are given by “Kernels” =(*k*1, *k*2, *k*3) and “Channels” =(*d*1, *d*2, *d*3) in Figure 3, and the shape of the input tensor is (d0,H,W).

**Figure 6:**
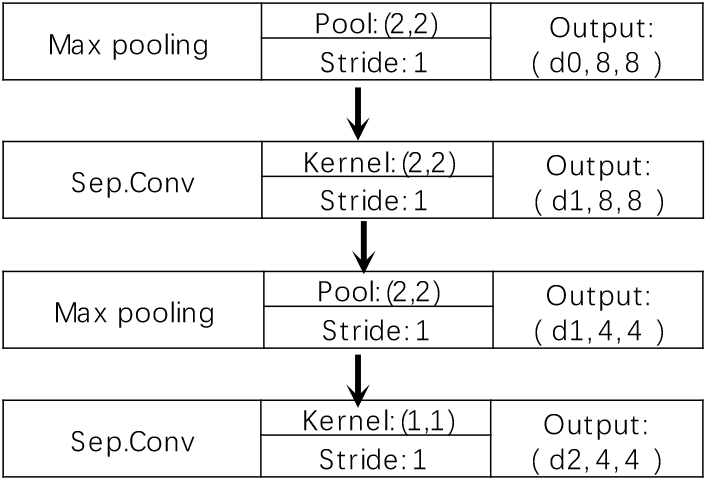
Network architecture of Block 3. Inputs are given by “Channels” = (*d*1, *d*2, *d*3) in Figure 3. Sep. Conv. = separable convolution layer (Gao *et al.*, 2018).

**Figure 7:**
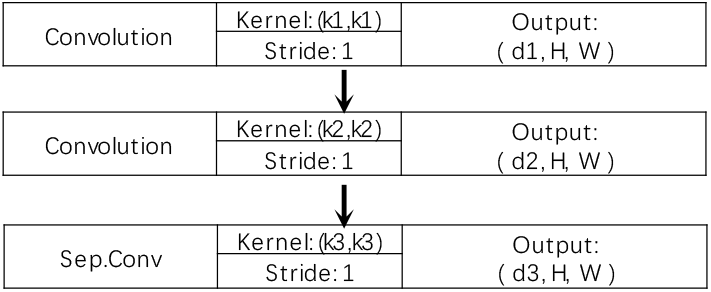
Network architecture of Block 4. Inputs are given by “Channels” = (*d*1, *d*2, *d*3) in Figure 3, and the shape of the input tensor is (d0,H,W). Sep. Conv. = separable convolution layer (Gao *et al.*, 2018).

## 4 Results

### 4.1 Single promoter classifiers select ALS-associated genes as well as potential novel risk factors

Figure 8 shows histograms of the classification accuracy for the single promoter classifiers, organized per chromosome. While most promoter regions lead to an accuracy around 0.5 - the same as random - the distribution has a tail on the right with a few promoter regions achieving higher accuracy. Hence only a few promoter regions have a potential relevance to ALS prevalence.

**Figure 8:**
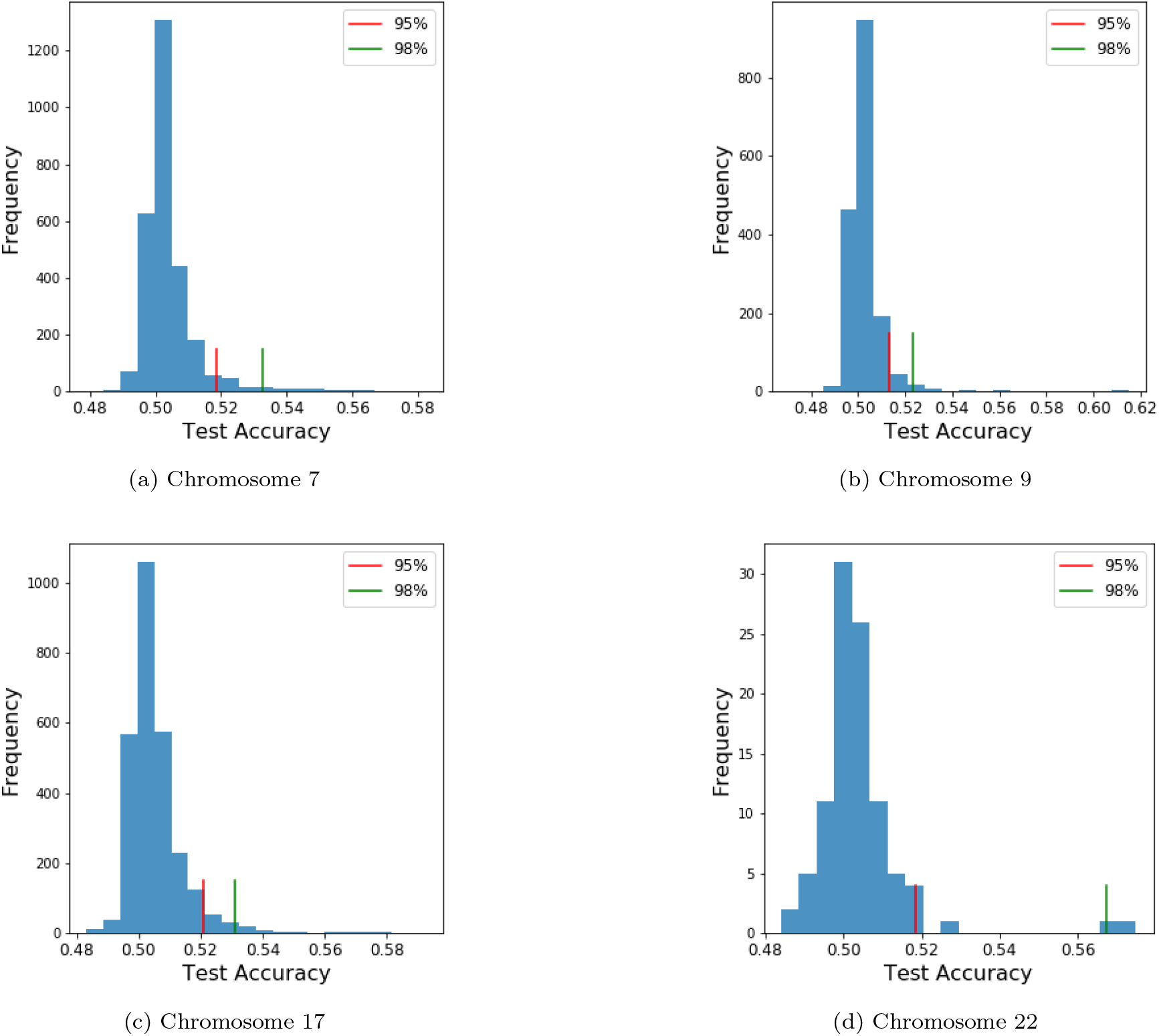
Histograms of per-promoter region test accuracies for each chromosome.

The genes that the selected promoter regions correspond to are listed in Table 2 together with the accuracy, precision and recall obtained with Promoter-CNN. Some of these genes have been associated with ALS or other neurological disorders before, while others can be viewed as potential novel ALS-associated genes. The accuracies for these promoter regions obtained by running a logistic regression are presented as well. The results show that using logistic regression would have resulted in a partially different selection of promoter regions. Recall that multiple promoter regions can correspond to a single gene, as a gene can have multiple transcription start sites.

**Table 2:** Promoter regions selected by the deep neural network for individual promoters. Accuracy (Acc), precision (Prec) and recall obtained with Promoter-CNN are reported. Additionally, the accuracy for this promoter region obtained with logistic regression is reported (Acc Log Regr). ^1^Reported as ALS associated gene (http://alsod.iop.kcl.ac.uk/) (Abel *et al.*, 2013). ^2^Reported to be associated with ALS and other neurodegenerative disorders (https://www.wikigenes.org/e/gene/e/2186.html).

|  | Positions (range) | Gene | Acc | Prec | Recall |
| --- | --- | --- | --- | --- | --- |
| Chr 7 | 72317552 - 72663411 | TYW1B | 0.566 | 0.672 | 0.259 |
|  | 76486174 - 76514143 | DTX2 | 0.562 | 0.719 | 0.210 |
|  | 108000837 - 108023643 | LAMB4 | 0.582 | 0.679 | 0.313 |
|  | 108001655 - 108023688 | LAMB4 | 0.576 | 0.668 | 0.304 |
|  | 142254448 - 142274025 | TRY2P | 0.567 | 0.717 | 0.222 |
|  | 60061 - 93116 | LOC105375113 | 0.579 | 0.728 | 0.252 |
|  | 68920 - 94119 | LOC101929756 | 0.566 | 0.672 | 0.259 |
|  | 157112225 - 157119549 | LOC105375607 | 0.576 | 0.745 | 0.230 |
| Chr 9 | 134626387 - 134643306 | COL5A1 | 0.600 | 0.710 | 0.338 |
|  | 136324634 - 136347520 | GPSM1 | 0.581 | 0.653 | 0.345 |
|  | 136334910 - 136355183 | GPSM1 | 0.615 | 0.694 | 0.412 |
|  | 136336933 - 136357677 | GPSM1 | 0.611 | 0.693 | 0.398 |
|  | 136361582 - 136378662 | SNAPC4 | 0.609 | 0.635 | 0.514 |
|  | 136370707 - 136389390 | SNAPC4 | 0.603 | 0.677 | 0.395 |
|  | 136382485 - 136403936 | SDCCAG3 | 0.581 | 0.673 | 0.314 |
|  | 98535635 - 98557461 | LOC105375972 | 0.595 | 0.753 | 0.282 |
| Chr 17 | 138726 - 158754 | DOC2B<sup>1</sup> | 0.582 | 0.603 | 0.477 |
|  | 139747 - 159851 | DOC2B<sup>1</sup> | 0.579 | 0.593 | 0.506 |
|  | 139853 - 160092 | DOC2B<sup>1</sup> | 0.573 | 0.601 | 0.434 |
|  | 141150 - 160523 | DOC2B<sup>1</sup> | 0.577 | 0.599 | 0.464 |
|  | 15614822 - 15661462 | TRIM16 | 0.591 | 0.793 | 0.250 |
|  | 55245746 - 55267188 | HLF | 0.577 | 0.747 | 0.234 |
|  | 55247456 - 55268383 | HLF | 0.577 | 0.745 | 0.230 |
|  | 67877749 - 67891928 | BPTF<sup>2</sup> | 0.592 | 0.793 | 0.250 |
| Chr 22 | 17230116 - 17293295 | CECR1 | 0.529 | 0.627 | 0.141 |
|  | 17541582 - 17566229 | CECR2 | 0.515 | 0.725 | 0.049 |
|  | 17747340 - 17783351 | BID | 0.518 | 0.598 | 0.113 |
|  | 17760620 - 17788896 | MIR3198-1 | 0.517 | 0.589 | 0.111 |
|  | 19149580 - 19162696 | GSC2 | 0.517 | 0.647 | 0.074 |
|  | 19403582 - 19439165 | HIRA | 0.5750 | 0.694 | 0.267 |
|  | 19404552 - 19439540 | HIRA | 0.567 | 0.694 | 0.267 |
|  | 19685895 - 19718733 | LINC00895 | 0.520 | 0.554 | 0.207 |

Several genes that were associated with ALS by earlier studies are not among the top eight performing promoter regions from Promoter-CNN. The classification accuracies from Promoter-CNN for the ALS-associated genes reported by Abel *et al.* (2013) are listed in Table 3. The ALS-associated genes from Abel *et al.* (2013) for chromosome 22 were not kept in our dataset after QC, and hence no results are reported.

**Table 3:** Accuracies for ALS-associated genes of chromosomes 7, 9, 17 and 22 as listed on http://alsod.iop.kcl.ac.uk/ (Abel *et al.*, 2013).

| Chr | Gene | Accuracy |
| --- | --- | --- |
| Chr7 | GARS | 0.5159 |
|  | RAMP3 | 0.503 |
|  | ZNF746 | 0.524 |
|  | DPP6 | 0.532 |
| Chr9 | SUSD1 | 0.504 |
|  | ALAD | 0.513 |
|  | STEX | 0.545 |
| Chr17 | MAPT | 0.514 |
|  | SLC39A11 | 0.511 |

### 4.2 ALS-Net outperforms other classifiers in terms of accuracy and recall

The selected promoter regions were included in a final overall classifier. We compared the performance of ALS-Net with logistic regression, SVM, random forest and AdaBoost (indicated by Promoter-CNN + classifier, Table 4). Additionally, we compare the results of Promoter-CNN with the five classifiers to logistic regression on all promoter regions, so without the help of Promoter-CNN. This was only possible for the individual chromosomes, as a logistic regression on all promoter regions from the four chromosomes combined required too much RAM. SVM, random forest and AdaBoost could not deal with the full chromosome data of even a single chromosome. The methods are compared based on classification accuracy, precision, recall and the F1 statistic for each chromosome separately as well as for their combination, and results are presented in Table 4.

**Table 4:** Classification results obtained with four classification methods applied to chromosomes 7, 9, 17 and 22 independently and combined. The result of best performing model for the given (set of) chromosome(s) is denoted in italic, while the overall best score is indicated in bold. Chr = chromosome.

| Classifier | Chr | Accuracy | Precision | Recall | F1-Score |
| --- | --- | --- | --- | --- | --- |
| Logistic Regression | 7 | 0.625 | 0.642 | *0.566* | 0.602 |
|  | 9 | 0.546 | 0.575 | 0.355 | 0.439 |
|  | 17 | 0.637 | 0.670 | 0.539 | 0.598 |
|  | 22 | 0.590 | 0.619 | *0.467* | *0.533* |
| Promoter-CNN + ALS-Net | 7 | *0.676* | 0.742 | 0.538 | *0.624* |
|  | 9 | *0.697* | 0.698 | *0.696* | *0.697* |
|  | 17 | *0.688* | 0.764 | *0.573* | *0.648* |
|  | 22 | *0.603* | *0.747* | 0.313 | 0.448 |
|  | All | **0.769** | 0.711 | **0.908** | **0.797** |
| Promoter-CNN + Logistic Regression | 7 | 0.635 | 0.728 | 0.445 | 0.553 |
|  | 9 | 0.683 | 0.743 | 0.560 | 0.685 |
|  | 17 | 0.642 | 0.734 | 0.445 | 0.554 |
|  | 22 | 0.580 | 0.714 | 0.299 | 0.422 |
|  | All | 0.739 | 0.759 | 0.699 | 0.728 |
| Promoter-CNN + SVM | 7 | 0.550 | 0.750 | 0.151 | 0.252 |
|  | 9 | 0.598 | *0.790* | 0.266 | 0.397 |
|  | 17 | 0.577 | *0.788* | 0.212 | 0.334 |
|  | 22 | 0.521 | 0.743 | 0.267 | 0.393 |
|  | All | 0.725 | 0.783 | 0.624 | 0.694 |
| Promoter-CNN + Random Forest | 7 | 0.562 | *0.776* | 0.175 | 0.285 |
|  | 9 | 0.579 | 0.759 | 0.229 | 0.351 |
|  | 17 | 0.645 | 0.762 | 0.420 | 0.542 |
|  | 22 | 0.587 | 0.745 | 0.265 | 0.391 |
|  | All | 0.596 | **0.813** | 0.249 | 0.381 |
| Promoter-CNN + AdaBoost | 7 | 0.604 | 0.642 | 0.467 | 0.541 |
|  | 9 | 0.621 | 0.668 | 0.481 | 0.559 |
|  | 17 | 0.599 | 0.633 | 0.472 | 0.401 |
|  | 22 | 0.561 | 0.591 | 0.398 | 0.475 |
|  | All | 0.661 | 0.700 | 0.565 | 0.625 |

First note that the classification accuracy of logistic regression is improved by Promoter-CNN. Second, Promoter-CNN + ALS-Net outperforms all other methods in terms of accuracy, closely followed by Promoter-CNN + logistic regression. Both methods largely outperform SVM, random forest and AdaBoost. Third, Promoter-CNN + ALS-Net almost always yields the highest recall, but is almost always outperformed by logistic regression, SVM and random forest in terms of precision – i.e., ALS-Net is better at identifying ALS patients (lower number of false negatives) but classifies healthy controls more often as patients than the other methods (higher number of false positives). Thus, each of the methods provides a different trade-off of precision versus recall. We therefore also consider the F1-statistic, a combined measure of precision and recall. Our deep neural network outperforms the other methods in terms of the F1 statistic for three out of the four individual chromosomes as well as the combination of chromosomes.

### 4.3 Including more genomic information improves classification

For most models the highest accuracy is obtained when the four chromosomes are combined rather than considering each chromosome individually. This does not hold for Promoter-CNN + random forest, which is likely due to the fact that a larger forest would be required to be able to deal with the larger dataset that is obtained when combining the three chromosomes.

### 4.4 ALS-Net is less sensitive to batch induced confounding effects

Genome sequences were obtained in 4 batches (C1, C3, C5, C44). Disease status is highly confounded with the batch an individual belongs to: the number of cases / controls was 226 / 380 for batch C1, 131 / 49 for batch C3, 0 / 5156 for batch C5 and 4154 / 1812 for batch C44. Together C44 and C5, two highly unbalanced batches, cover approximately 93% of the individuals. A classifier may thus achieve good accuracy on predicting disease status by picking up batch-related data structures rather than disease-associated genetic characteristics. If a classifier picks up differences between C5 and C44, instead of between case (ALS) and control (no ALS), the classifier will fail to make reasonable predictions in C1 and C3, which still cover 786 individuals. Since both C1 and C3 are fairly balanced in terms of case-control labels, batch labels cannot be confounded with true case / control labels as easily as in C5 and C44. To check whether Promoter-CNN + ALS-Net (our approach) and Promoter-CNN + logistic regression (as the second best classifier evaluated) pick up on disease status rather than batch effects during training we evaluated the performance of these two within the individual batches on the training data. As can be seen in Table 5 both classifiers achieve good accuracy within batches C5 and C44, for which it remains unclear whether the classifiers predict batch labels rather than true labels. On C1 and C3 Promoter-CNN + logistic regression fails to bring up competitive performance rates (in particular: precision/recall logistic regression: 0.48 / 0.37 on C1 and 0.77 / 0.31 on C3), while Promoter-CNN + ALS-Net keeps significantly better performance rates (precision/recall: 0.51 / 0.79 on C1, 0.83 / 0.76 on C3), a clear indication that the predictions of ALS-Net are due to picking up truly ALS related effects to a substantial amount. For logistic regression however, it is likely that batch effects have been picked up, which lead to random classification in batches C1 and C3.

**Table 5:**
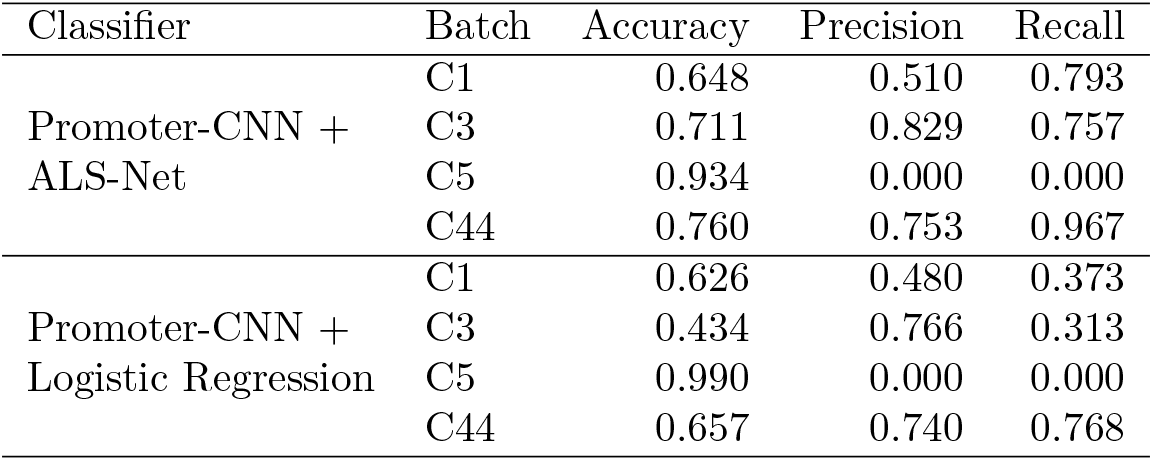
Training accuracy, precision and recall for each cohort, obtained with Promoter-CNN + Logistic regression and Promoter-CNN + ALS-Net.

While ALS-Net comes with the clear promise to be (considerably) less prone to picking up batch effects, we conclude to say that correcting for confounding effects for CNN based methods still requires further research, which we consider most interesting future work.

### 4.5 Validation by testing on other phenotypes

Although the network architecture was optimized for chromosome 7, no adjustments needed to be made for the other three chromosomes and their combination. This suggests that our network architecture generalizes well. We therefore wish to test whether the model is specific for ALS classification, or also classifies well for other sample characteristics that are potentially confounded with disease status. We do so by using the optimized model architecture and train it for the prediction of gender. The model achieved a test accuracy of only 52.27%, hardly better than random. This observation supports the hypothesis that our network architecture is specific to ALS classification.

### 4.6 Runtimes of ALS-Net are acceptable

Promoter-CNN was run on a CPU cluster. The training process for a single promoter region takes around 200s. Note that training of the promoter regions can be done in parallel on a multi-processor system. ALS-Net was trained on a GPU (Nvidia TitanX). For the largest model, which classified individuals based on information from the four chromosomes combined, the model needed 500s for training.

## 5 Discussion

In this work we presented a novel deep learning-based approach for genotype-phenotype association studies on genome-sized data that allows for the identification of phenotype-associated genes. We used the earlier observed hypersensitivity of regulatory elements on the genome for developing a two-step approach, and designed a neural network where we made use of the structure of genome data by first considering each promoter region separately, and then combining their information in later layers. The combination of Promoter-CNN with ALS-Net has several advantages: it (1) can handle genome-sized data, (2) identifies regions of the genome that are relevant to classification of ALS patients versus healthy controls, and (3) yields good classification results.

ALS-Net generally outperforms other methods in terms of classification accuracy, followed by Promoter-CNN aided logistic regression. As for prior related work, note that Bellot *et al.* (2018) observed small improvements of CNN based methods over logistic regression in several, but not all cases. Here we observe some marked improvements of Promoter-CNN + ALS-Net over logistic regression. An explanation might be that Bellot *et al.* (2018) use a (substantially) simpler (nondeep) network architecture and make a pre-selection of genetic features based on linear models, and hence overlook non-additive interactions already in the pre-selection step.

ALS-Net outperforms all other methods in terms of recall, also called power. This indicates that our approach might point out ways to overcome the (notoriously complained) lack of power that arises from the use of linear models when associating genotypes with phenotypes that underlie more involved genetic architectures. Additionally further examination of what caused the increase in true case predictions might yield novel insight in the genomic mechanisms underlying ALS. Overall, ALS-Net provides a better trade-off between precision and recall as measured by the F1 statistic, which finally documents its value as a predictor in general.

Note that all methods have been helped by our two-step approach: the classification performance of logistic regression goes up when combined with Promoter-CNN, while none of the other methods evaluated were able to process chromosome-sized genotype data without pre-selecting features.

Our results support the belief that ALS is caused by non-linear combinations of variants, which was hypothesized before by Van Rheenen *et al.* (2016). Table 4 shows a low recall for each of the individual promoter regions. Combining these into a single classifier improves recall for PromoterCNN + ALS-Net, while this effect is much smaller for logistic regression as well as PromoterCNN + logistic regression. This implies that the promoter regions on the different chromosomes interact in a non-additive way.

Our analysis has identified several promoter regions that potentially contribute to ALS prevalence, some of which are known to be associated with ALS. On the other hand, several ALS-associated genes (Abel *et al.*, 2013) were not selected by Promoter-CNN. This does not necessarily imply that Promoter-CNN gave low prediction accuracies for these promoter regions: they simply were not among the eight most predictive promoter regions. Four out of the nine ALS-associated promoter regions that were not selected by Promoter-CNN were among the 5% best performing promoter regions (for these promoter regions, Promoter-CNN achieved an accuracy above 0.518, 0.513 and 0.520 for chromosomes 7, 9 and 17 respectively, see Figure 8). Despite missing some of the known ALS-associated genes, our final classification, which did not use any information from these genes, was able to classify at high accuracy. Further research is required to understand this.

The architecture of ALS-Net was optimized for chromosome 7. When applying this architecture to classify samples from the test set based on genotype data from chromosomes 9, 17 and 22 as well as their combination, the model performed very well and there was no need for further adjustment of the network architecture. These results show that our network architecture generalizes well to unseen data, even to data from a different chromosome or set of chromosomes. Since beyond generally applicable genetics principles we have not made use of particular ALS related knowledge, we believe that our architectures hold the potential to be applicable more universally. Further such experiments, however, predominantly depend on the availability of cohorts of sizes equal to the rather large cohorts we have been investigating here – which will be possible for ever more diseases in the mid-term future.

A major issue for genotype-phenotype association studies has been the large number of input variables, which causes issues for most machine learning approaches. To the best of our knowledge, there has so far been no method that can deal with more than half a million of genetic variants other than GWAS (where one tests for the association of a single variant with genotype) or approaches where GWAS or prior knowledge was used for pre-selecting relevant variants. This makes our approach the first that accounts for non-additive interactions between genomic features right from the start.

We view our work as a first step towards biology-informed deep learning for association studies. While our results are promising, several improvements can still be made. First, we chose to select the top eight performing promoter regions. Including more promoter regions may improve performance, but will also lead to an increase in computational burden. Also the number of included promoter regions may be chosen to be dependent on the length or the expected contribution to heritability of the chromosome under consideration. Additionally, an even deeper model may improve performance as well. We plan to further develop these methods in the future.

While this work presents a methodology for the analysis of genotype-phenotype data, refinements are required before practical implementation. For example, in our analysis we did not account for population stratification. As we first focus on the development of the neural network-based approach, we leave such improvements for future research.

Even though our analysis was limited to the genomic information of only four chromosomes, we obtained a high level of classification accuracy. We plan to extend our work by including all chromosomes, which we expect to result in a strong increase in recall and hence classification accuracy, as well as the identification of more potential ALS-associated promoter regions. Additionally, ProjectMinE is a worldwide ongoing effort, and we plan to apply our methods to the full dataset to strengthen our results once this data becomes available.

## 6 Conclusion

In this paper we presented ALS-Net, a convolutional neural network approach to predict ALS prevalence from genotype data. In order to employ the strengths of convolution we have developed a two-level approach where we focus on promoter regions, which are known sensitive sites for disease-causing variants. The architecture of the final classification network employs the strength of convolution and the structure of genome data by applying convolution filters to individual promoter regions. The results of our tests are promising, and are expected to generalize to genome regions that were unexplored in this work. Additionally, this work shows that deep learning is a highly promising approach for the identification of complex genotype-disease relations. We view our approach as a first step towards deep learning for genotype-phenotype association analysis guided by regulatory principles.

